# Decoding *Helicobacter pylori* Resistance: Machine Learning–Enhanced Prediction of Antibiotic Susceptibility using Whole-Genome Sequencing

**DOI:** 10.1101/2025.08.07.669068

**Authors:** Florent Ailloud, Diyuan Lu, Beate Spiessberger, Daksh Pamar, Atefeh Kazeroonian, Michael Flossdorf, Monica Oleastro, Christian Schulz, Markus Gerhard, Michael P. Menden, Sebastian Suerbaum

## Abstract

**Background:** *Helicobacter pylori* is a significant risk factor for gastric cancer, peptic ulcers, and MALT lymphoma. Rising antibiotic resistance rates complicate treatment strategies. While nucleotide sequence based assays are reliable in predicting clarithromycin and levofloxacin resistance, predicting metronidazole resistance is more challenging due to diverse metabolic pathways contributing to resistance, and high genomic variability.

**Methods:** We assembled a cohort of 483 *H. pylori* clinical isolates, combining whole-genome sequencing with phenotypic susceptibility testing. Machine learning models (SVM, XGBoost, FNN) were trained on genomic variants to predict resistance phenotypes. A sliding-window approach and SHAP-based importance scoring were used for feature selection to identify biologically relevant mutations, improving prediction accuracy, particularly for metronidazole resistance.

**Results:** The best-performing FNN model improved metronidazole resistance prediction by 16% compared to conventional (non-ML, single polymorphisms) sequence-based detection methods applied to the same strain collection. Feature selection identified 32 feature sets, with 11 sets significantly improving F1-scores over the baseline. Combining 2–4 feature sets revealed 53 synergistic combinations across all models. Validation showed that 87% of these combinations significantly outperformed non-ML molecular testing, with 16 combinations achieving F1-scores above 0.65.

**Conclusion:** Machine-learning can significantly improve the performance of sequence-based susceptibility testing for metronidazole in *H. pylori*. Novel candidate predictive markers identified from whole-genome data offer testable hypotheses about yet unexplored mechanisms of metronidazole resistance. These findings support the potential for ML-based approaches to enable more accurate susceptibility-guided therapies.

## 1 Introduction

*Helicobacter pylori* (*H. pylori*) infects approximately half of the global population and is a major risk factor for gastric cancer, as well as a leading cause of peptic ulcer disease and gastric MALT lymphoma.^1^ Typically acquired in childhood through person-to-person transmission, the infection often persists throughout life if left untreated. Screening strategies for *H. pylori* and gastric cancer are implemented in high prevalence regions, mostly in Asia.^2,3^ Despite these strategies, an increased number of gastric cancer cases attributable to *H. pylori* is expected for young people born in 2008-2017^4^.

Current management of *H. pylori* infection relies exclusively on antibiotic-based combination therapies; a vaccine is not available despite decades of research. First-line eradication regimens include triple or quadruple therapies combining a proton pump inhibitor with two antibiotics (and bismuth in the case of quadruple therapy). Antibiotics widely used in such regimens include clarithromycin, metronidazole, amoxicillin, tetracycline, or levofloxacin.^1^ If successful, eradication of *H. pylori* significantly reduces gastric cancer incidence^5^, heals recurrent ulcer disease, and induces complete remissions of early MALT lymphomas in a large number of cases. Antibiotic resistance is considered a critical factor in treatment failure.^6,7^ Despite this, access to antibiotic susceptibility testing remains limited for availability and reimbursement reasons from a global perspective.^8^

In recent years, machine learning (ML) has been applied in the *H. pylori* field, to improve detection via endoscopy and histopathology.^9–12^ Due to the high genetic diversity of *H. pylori*^13^ and the complexity of some of its antibiotic resistance mechanisms^14^, ML represents a promising approach to investigate genotype-phenotype associations and establish predictive models of antibiotic susceptibility in this pathogen. However, a major limitation to developing such models has been the scarcity of datasets from *H. pylori* clinical isolates that have been characterized both genetically and phenotypically for resistance to relevant antibiotics. In this study, we assembled a large-scale discovery cohort of 483 *H. pylori* clinical isolates, each characterized by whole-genome sequencing and culture-based antibiotic susceptibility testing for clarithromycin, levofloxacin, and metronidazole. Using this dataset, we developed a feature selection pipeline based on genomic coordinates to generate sets of polymorphic variants suitable for training machine learning and deep learning models, with the aim to predict resistance phenotypes. An independent validation cohort of 50 additional clinical isolates, used solely for external testing, confirmed the generalizability of our models. The best-performing models improved the accuracy of metronidazole resistance prediction by 16% compared to non-ML sequence-based diagnostic approaches. This dataset further enables future investigations into the genetic basis of *H. pylori* drug resistance.

## 2. Results

### 2.1 Training dataset assembly and identification of genome-wide variants in *H. pylori*

To train ML models for antimicrobial resistance prediction, we assembled a collection of 483 *H. pylori* strains from three different laboratories (referred to as the HpRES-500 dataset). All strains were recovered initially from patient gastric biopsy specimens during routine endoscopy. The largest group consists of 222 clinical isolates obtained from the National Reference Center for *H. pylori* in Munich (Max von Pettenkofer Institute, Germany), followed by 152 strains belonging to the laboratory collection of the Institute of Medical Microbiology at the Technical University of Munich (Germany) and 109 clinical isolates from the National Reference Center for *H. pylori* in Lisbon (Portugal).

Phenotypic resistance data obtained via accredited culture-based methods were collected for clarithromycin, levofloxacin and metronidazole. To reduce class imbalance and support reliable model training, both resistant and susceptible isolates were selectively included across antibiotic classes. As a result, the dataset maintains a balanced distribution, with each antibiotic represented by at least 20 percent resistant strains (Supplementary Table S1).

In the context of training classification models, phenotypic resistance data represent the output variables (or targets) of the ML algorithms. To generate the input variables (or features), we performed whole-genome sequencing of *H. pylori* strains and identified an average of 67,465 ± 7,901 genotypic variants per isolate, resulting in a total of over 30 million variants across all isolates. Because the number of variants is substantially larger than the number of strains, this dataset would typically be considered high-dimensional, which increases the computational cost and the risk of overfitting (i.e., ML models capturing random noise rather than the underlying relationships within the dataset).

To retain the most important biological context while decreasing the number of features, variants were filtered and grouped according to their predicted functional effects (see *Methods*): non-coding variants (Category A), amino acid substitutions (Category B), and loss-of-function mutations (Category C). This first feature reduction step allowed us to compress our dataset by 70%. Accordingly, all known biomarkers associated with drug resistance in *H. pylori* fall into one of these categories, suggesting that our feature set covers a wide range of potential resistance mechanisms.

### 2.2 Performance of established (single polymorphism, non-ML) nucleotide sequence-based susceptibility testing

The susceptibility of *H. pylori* to clarithromycin and levofloxacin can be reliably predicted by detecting high-confidence genetic polymorphisms using diverse molecular methods (qPCR, Sanger sequencing, LAMP, targeted sequencing), which have been implemented in several commercially available diagnostic kits.^15,16, 17,18^ In contrast, susceptibility to metronidazole cannot be reliably detected using known genetic markers, and only experimental markers have been reported.^19^ To establish a baseline for the evaluation of ML performance, we selected the most frequently used and best performing genetic markers included in commercial methods or in the literature (Supplementary table S2) to predict drug susceptibility in the discovery collection of *H. pylori* strains (HpRES-500).

Sequence-based molecular predictions achieved high accuracy, specificity, and sensitivity for clarithromycin and levofloxacin (Table 1), while performance for metronidazole was substantially lower. In particular, the reduced sensitivity indicates poor detection of resistant isolates, leading to many false negatives. Low F1-score and Matthews correlation coefficient (MCC) further confirm that conventional sequence-based assays are insufficient for reliably predicting metronidazole resistance in *H. pylori*.

**Table 1.** Performance of univariate genetic resistance markers (Supplementary Table S2) using biostatistical evaluation. Predictive performance of established individual resistance-associated variants evaluated in the *HpRES-500* dataset using standard biostatistical metrics.

| Antibiotic | Acc. (%) | Sens. (%) | Spec. (%) | F1-score (%) | MCC |
| --- | --- | --- | --- | --- | --- |
| Clarithromycin | 92.3 | 84.7 | 97.1 | 89.5 | 0.84 |
| Levofloxacin | 90.7 | 73.1 | 98.5 | 82.9 | 0.78 |
| Metronidazole | 65.2 | 40.2 | 97.0 | 56.4 | 0.43 |

### 2.3 Machine learning performance utilising known resistance markers

Classic sequence-based susceptibility testing is limited by lacks of any decision algorithm and typically classifies a sample as resistant if any known resistance marker is detected. This approach requires high-confidence markers to balance sensitivity and specificity. In contrast, ML can potentially capture the relative contribution of each marker during training and integrate all available information into a classification model, including low-confidence markers with limited or no experimental validation.

We trained support vector machines (SVM), extreme gradient boosting (XGBoost), and feedforward neural networks (FNN; Methods) using all known resistance markers ^14^. All models showed improved performance, with high accuracy for clarithromycin (⍰F1 = 3%; AUC = 90.1–91.0) and levofloxacin (up to ⍰F1 = 8.5%; AUC = 86.7–90.6), shown in Table 2. Metronidazole predictions remained variable but improved (up to ⍰F1 = 8.7%; AUC = 53.7–75.0), underscoring the benefits of multivariate ML over traditional univariate sequence-based methods. Notably, ML models improved the sensitivity of antibiotic resistance prediction relative to established sequence-based molecular testing. Sensitivity increased by approximately 5% for clarithromycin and 15% for levofloxacin. For metronidazole, sensitivity improved by up to 25%, suggesting that ML approaches more effectively identified true resistant cases.

**Table 2.** Performance of ML models trained on known sequence-based resistance markers. Predictive accuracy, F1-score, MCC, sensitivity, specificity, and AUC values are reported for SVM, XGBoost, and FNN classifiers trained on known resistance-associated variants in the *HpRES-500* dataset.

| Antibiotic | ML | AUC | Acc. (%) | Sen. (%) | Spe. (%) | F1score (%) | MCC |
| --- | --- | --- | --- | --- | --- | --- | --- |
| Clarithromycin | SVM | 90.5 ± 0.2 | 93.0 ± 0.0 | 91.8 ± 0.0 | 97.1 ± 0.0 | 92.5 ± 0.0 | 0.85 ± 0.00 |
|  | XGBoost | 90.1 ± 0.6 | 93.0 ± 0.0 | 91.8 ± 0.0 | 97.1 ± 0.0 | 92.5 ± 0.0 | 0.85 ± 0.00 |
|  | FNN | 91.0 ± 0.7 | 93.0 ± 0.0 | 91.8 ± 0.0 | 97.1 ± 0.0 | 92.5 ± 0.0 | 0.85 ± 0.00 |
| Levofloxacin | SVM | 89.0 ± 0.8 | 92.8 ± 0.1 | 90.5 ± 0.2 | 96.5 ± 0.2 | 91.4 ± 0.1 | 0.83 ± 0.00 |
|  | XGBoost | 86.7 ± 0.6 | 92.0 ± 0.0 | 89.1 ± 0.0 | 96.6 ± 0.0 | 90.3 ± 0.0 | 0.81 ± 0.00 |
|  | FNN | 90.6 ± 0.5 | 92.6 ± 0.3 | 89.8 ± 0.8 | 97.1 ± 0.5 | 91.0 ± 0.5 | 0.82 ± 0.01 |
| Metronidazole | SVM | 53.7 ± 1.8 | 56.0 ± 1.6 | 53.7 ± 1.8 | 11.2 ± 5.5 | 44.3 ± 4.7 | 0.13 ± 0.06 |
|  | XGBoost | 65.1 ± 1.5 | 65.3 ± 1.3 | 65.1 ± 1.5 | 62.3 ± 4.4 | 65.1 ± 1.5 | 0.30 ± 0.03 |
|  | FNN | 75.0 ± 0.3 | 67.2 ± 0.5 | 68.7 ± 0.5 | 97.5 ± 0.6 | 65.0 ± 0.7 | 0.45 ± 0.01 |

Clarithromycin and levofloxacin showed consistent performance across all three ML algorithms. In contrast, for metronidazole, the FNN yielded the highest overall performance, outperforming established sequence-based testing across all evaluated metrics. These differences highlight how each algorithm handles underlying data structure and class imbalance, with FNN particularly benefiting from its capacity to model non-linear relationships. For metronidazole, the feedforward neural network (FNN) improved the F1-score by 10% over non-ML, single polymorphism-based testing, with similar accuracy and MCC. Sensitivity increased by 28% but remained below 70%, suggesting that existing markers do not capture all resistance cases. To identify potentially novel genetic markers associated with resistance to metronidazole in a broader range of isolates, ML can be applied to genetic variants representing the whole genome of *H. pylori*.

### 2.4 Genome-Wide Identification of Drug Resistance Markers through Feature Ranking

High-dimensional genomic data pose a significant challenge for antibiotic resistance prediction in *H. pylori* using ML. In the present study, over 500,000 binary variant features were derived from 483 discovery isolates, resulting in a feature-to-sample ratio exceeding 1,000:1. Effective feature selection is therefore critical to reduce dimensionality, prevent overfitting, and retain biologically relevant variants, which range from single-nucleotide polymorphisms to loss-of-function mutations with varying predictive power.

Known mutations associated with antibiotic resistance are typically clustered within a limited number of genes. For example, 50 genetic variants connected to levofloxacin resistance are found within less than 500 bp in the gyrase A gene (*gyrA*). Consequently, we decided to use a sliding-window approach to partition the genome into 830 non-overlapping 2,000 bp windows and focused on neighboring features across small genomic regions. Each segment contained a median of 600 features, balancing resolution and computational efficiency.

To identify windows with informative features, we trained an ML model (XGboost) separately on each segment using a five-fold cross-validation strategy (see *Methods*). The training was performed for the prediction of metronidazole resistance as a proof-of-principle due to the lack of reliable molecular markers. Genomic windows with AUC ≥ 0.55 were deemed informative (Methods), identifying 11 regions with 6,807 features. To further reduce dimensionality, the discriminative power of each feature within each genomic window was determined by calculating feature importance scores (based on the Shapley analysis; see *Methods*). Using a feature importance threshold of 0.01, a total of 293 variants was selected and combined into 32 genomic clusters spanning up to 10kb (Supplementary Table S3).

### 2.5 Prediction of antibiotic resistance with supervised learning algorithms

Support vector machines (SVM), extreme gradient boosting (XGBoost), and feedforward neural networks (FNN) were trained to predict resistance to metronidazole using top-ranked features derived from the sliding-window and feature importance analysis. Antibiotic resistance was modeled as a binary classification task, where class 0 corresponds to susceptibility and class 1 to resistance, and performance was evaluated using the F1-score to account for class imbalance. To obtain robust performance estimates, we employed three evaluation strategies: a) averaging across multiple runs (Avr.), b) within-model aggregation (Agg.), and c) an ensemble combining predictions from all models and runs (Ensemble).

Focusing on metronidazole resistance, we evaluated the benefit of whole-genome features using the FNN model trained on known biomarkers as a baseline. Each of the 32 selected genomic windows was combined with these markers to train SVM, XGBoost, and FNN models. FNN showed the highest average F1-score (64%), followed by XGBoost (61%) and SVM (54%). Aggregated predictions improved scores by 1.84% and enhanced model robustness. FNN and Ensemble consistently outperformed SVM, with Ensemble improving performance stability and generalizability.

A subset of genomic feature sets significantly enhanced metronidazole resistance prediction beyond the baseline biomarker model. Specifically, 34% (11/32) of the feature sets showed significantly improved F1-scores (p *<* 0.05, BH-corrected Mann–Whitney U test; Cohen’s d *>* 0.8) in at least one ML model and evaluation strategy (Figure 1; Supplementary Table S4). Gains ranged from 0.90% to 12.58%, with a mean improvement of 3.29%. These informative variants were broadly distributed across the *H. pylori* genome and enriched in genes linked to cell wall, membrane, and envelope biogenesis (Supplementary Table S5).

**Fig. 1.**
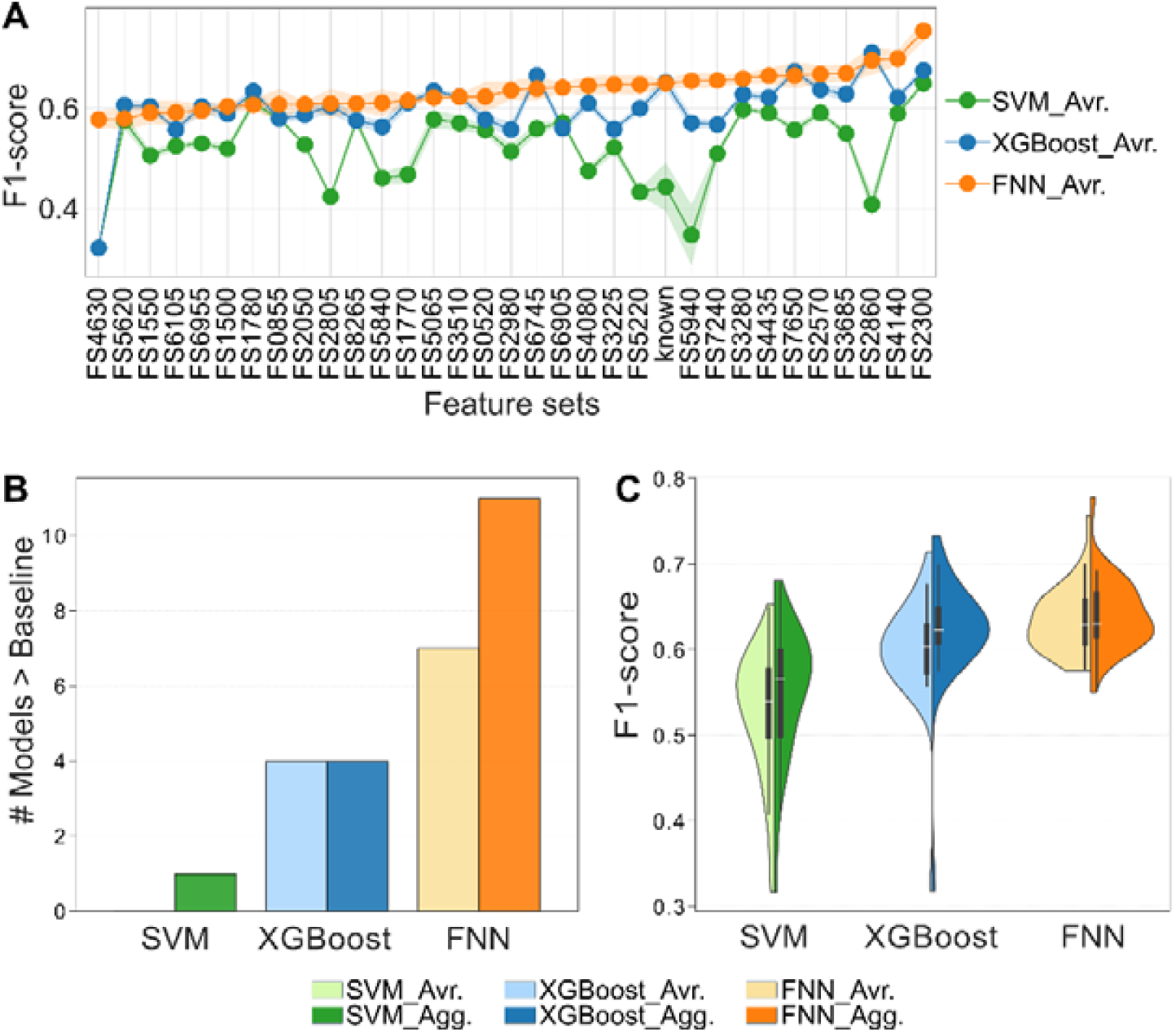
Performance comparison of different ML models for predicting drug resistance based on neighboring loci in close vicinity for Metronidazole in the withheld dataset. (A) F1-score performance across different single loci for three models: SVM, XGBoost, and FNN. The shaded region represents variance. (B) Count of significantly better-than-baseline predictive models for each ML method, in average (Avr.) and aggregate (Agg.) cases. (C) Violin plot of F1-scores for each model, comparing average (Avr.) and aggregate (Agg.) performance distributions.

### 2.6 Combinatorial Feature Optimization for Metronidazole Resistance Prediction

We investigated potential synergistic interactions among predictive feature sets to reflect the multifactorial nature of antibiotic resistance. ML models were trained across all possible combinations of the 11 previously identified feature sets that significantly outperformed the baseline, yielding over 2,000 subsets (six independent five-fold runs). A subset was defined as synergistic if its F1-score (i) significantly exceeded the baseline, (ii) demonstrated a large effect size (Cohen’s d *>* 0.8), and (iii) outperformed each individual constituent feature set (p *<* 0.05, BH-corrected Mann–Whitney U test; Methods).

Combinatorial feature sets enhanced the predictive performance. Out of 339 tested combinations, 53 met all criteria for synergy based on aggregated performance metrics, combining between two and four feature sets (Figure 2; Supplementary Table S4). Models with five or more sets failed to meet synergistic thresholds, indicating diminishing returns with increased feature complexity (Figure 5B). Pairwise combinations yielded the highest number of synergistic models (Figure 5A), suggesting that compact subsets of complementary features are most effective.

**Fig. 2.**
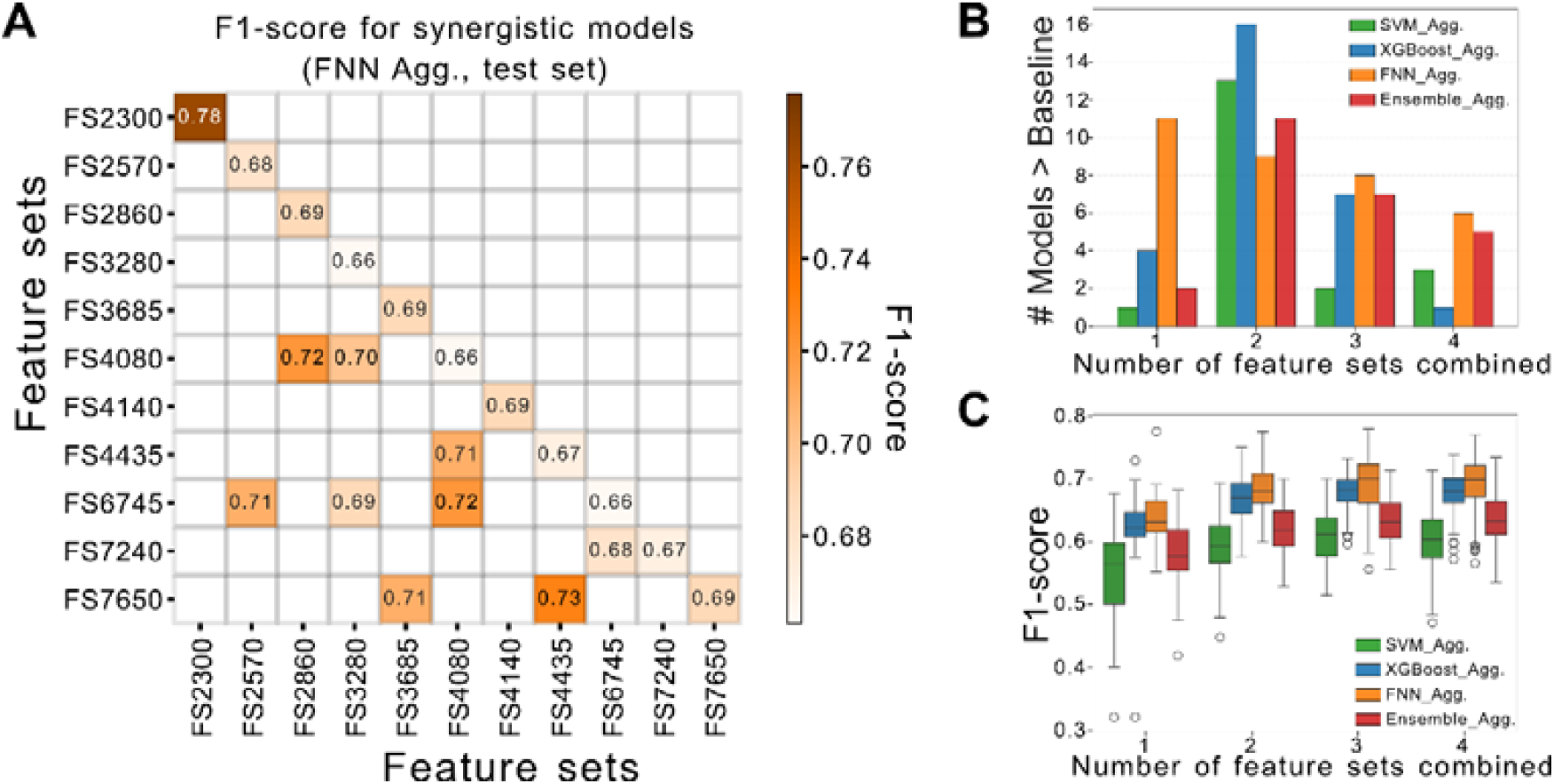
Synergistic feature set combinations enhance metronidazole resistance prediction in HpRES-500. (A) Heatmap of F1-scores for pairwise feature set combinations using FNN (aggregated). Diagonal shows individual set performance. Only combinations significantly exceeding baseline and components are shown (p < 0.05, BH-corrected; Cohen’s d > 0.8). (B) Number of significantly improved models by combination size across FNN, SVM, and XGBoost. (C) F1-score distributions by combination size and ML algorithm. Dashed line indicates baseline F1-score from established biomarkers.

Synergistic improvements in prediction performance vary by model type and feature contribution. Synergistic models were most frequently obtained using FNN and XGBoost (both 23/53), followed by Ensemble (16/53) and SVM (6/53). On average, these models improved the F1-score by 5% [range: 0.7–10%] relative to the baseline (Figure 5C). All 11 informative feature sets contributed to at least one synergistic combination, with FS2570 appearing most frequently (26/53 models). These findings demonstrate that selective combinations of genomic regions can substantially improve metronidazole resistance prediction beyond the use of individual sets alone.

The top-performing synergistic model leveraged complementary sequence-based genomic signals to achieve optimal predictive accuracy. This FNN-based model, comprising FS2570, FS3280, FS4435, and FS7650, yielded the highest F1-score (75%) and the largest gain over the baseline (10%). Compared to its individual components, the combination improved the F1-score by 7–9%, demonstrating additive predictive value. The involved genes spanned distinct COG functional categories, reinforcing the notion that metronidazole resistance in *H. pylori* arises from diverse and potentially complementary biological pathways (Supplementary Table S5).

### 2.7 Model Validation Using an Unseen Cohort of Clinical *H. pylori* Isolates

Validation on independent clinical isolates provides a critical test of model generalizability beyond the discovery dataset. We assembled a validation cohort of 50 *H. pylori* isolates from the National Reference Center for *H. pylori* (Germany), used exclusively for testing. These isolates were sequenced and phenotyped using protocols identical to the discovery cohort. Metronidazole resistance in the validation cohort was 58%, comparable to 56% in the discovery dataset. We first evaluated univariate biomarker-based predictions. Due to *H. pylori*’s extensive genomic diversity, established molecular testing showed reduced performance for metronidazole, with the F1-score decreasing from 56.4% in the discovery cohort to 33% in the validation set (Table 1; Table 3). In contrast, performance for clarithromycin and levofloxacin remained stable. These results underscore the limitations of fixed molecular diagnostics for metronidazole resistance and the need for improved models that accommodate more complex causation of resistance.

**Table 3.** Validation of univariate resistance markers using <u>biostatistical</u> model. Predictive performance of established resistance biomarkers in molecular susceptibility testing in the validation cohort.

| Antibiotic | Acc. (%) | Sen. (%) | Spec. (%) | F1-score (%) | MCC |
| --- | --- | --- | --- | --- | --- |
| Clarithromycin | 96.0<br>(86.3, 99.5) | 80.0<br>(44.4, 97.5) | 100.0<br>(91.2, 100.0) | 88.9<br>(66.7, 100.0) | 0.87<br>(0.67, 1.00) |
| Levofloxacin | 94.0<br>(83.5, 98.7) | 75.0<br>(42.8, 94.5) | 100.0<br>(90.7, 100.0) | 85.7<br>(66.7, 100.0) | 0.83<br>(0.66, 1.00) |
| Metronidazole | 52.0<br>(37.4, 66.3) | 20.7<br>(8.0, 39.7) | 95.2<br>(76.2, 99.9) | 33.3<br>(12.9, 51.6) | 0.23<br>(-0.03, 0.42) |

We next evaluated ML models trained exclusively on known metronidazole resistance biomarkers. While SVM maintained comparable F1-scores around 45% between the training and validation cohorts (Table 2; Table 4), this overall predictive performance remained low. In contrast, XGBoost and FNN models reduced F1 scores by 25% and 17%, respectively, in the validation cohort, indicating limited ability to generalize and potential overfitting. All models showed decreased sensitivity, reflecting reduced detection of resistant isolates. Despite this decline, the FNN model achieved the highest F1-score (48%; Table 4) and was used as the baseline for further validation analyses.

**Table 4.** Validation of ML models utilizing established resistance biomarkers for Metronidazole. Predictive performance of ML models trained with known metronidazole resistance biomarkers in <u>validation</u> cohort (averaged within models).

| ML | AUC | Acc. (%) | Sen. (%) | Spec. (%) | F1-score (%) | MCC |
| --- | --- | --- | --- | --- | --- | --- |
| SVM | 53.1 ± 1.5 | 57.1 ± 6.7 | 53.1 ± 1.5 | 28.2 ± 32.3 | 44.7 ± 7.1 | 0.11 ± 0.05 |
| XGBoost | 49.7 ± 2.1 | 47.0 ± 5.1 | 50.8 ± 1.2 | 74.4 ± 25.6 | 40.5 ± 10.9 | 0.02 ± 0.02 |
| FNN | 54.5 ± 1.0 | 52.1 ± 0.3 | 58.0 ± 0.2 | 95.2 ± 0.0 | 48.0 ± 0.4 | 0.23 ± 0.01 |

A majority of feature sets selected from the discovery cohort were successfully validated in the independent validation cohort. Specifically, 87% of the tested models (304/350) showed a significant F1-score improvement over the baseline (p *<* 0.05, BH-corrected Mann–Whitney U test; Cohen’s d *>* 0.8, based on aggregated performance). Validated models achieved an average F1-score gain of 4.3% [range: 0.6–13.0%] in the validation cohort and 8.5% [range: 0.7–24.4%] in the discovery cohort (Supplementary Table S4). FNN (n = 201) and XGBoost (n = 234) produced more validated models than Ensemble (n = 88) or SVM (n = 34). Notably, only 139 feature sets overlapped between FNN- and XGBoost-validated models, highlighting their complementary ability to extract diverse predictive signals. Among the validated feature sets, 16 achieved F1-scores above 0.65 with FNN in both discovery and validation cohorts, indicating consistent performance (Supplementary Table S6). One model combining FS2300, FS2570, FS4140, and FS4435 reached F1-scores of 0.73 and 0.72 in the discovery and validation datasets, respectively. Compared to classic molecular testing, it improved accuracy by 9–23%, mainly through a 33–51% gain in sensitivity, providing a more balanced prediction (Supplementary Table S7). These findings support the utility of selected genomic feature combinations for more robust resistance prediction.

### 2.8 Functional significance of validated feature sets associated with metronidazole resistance

The functional basis of metronidazole resistance in *H. pylori* is not completely understood and lacks support from both experimental and clinical data. The best characterized mechanism involves inactivation or mutation of the NAD(P)H nitroreductase RdxA, which plays a key role in the drug’s reductive activation and generation of cytotoxic radicals.^14,20^ Additional mechanisms may involve mutations in DNA repair and oxidative stress response genes, or changes in membrane proteins affecting drug uptake or efflux. To evaluate the contribution of known and newly predicted resistance markers within each ML model, we calculated feature importance scores (SHAP) for each genetic variant in our validated sets.^21^ In the best-performing FNN model trained only on known sequence-based markers (Table 2), *rdxA* and *frxA* loss-of-function variants, along with RdxA R16 mutations, were the strongest contributors. Mutations that lead to RecA H103 and RibF T222 amino acid exchanges also showed notable importance.

For models trained on newly predicted sequence-based markers, we selected genotypes based on experimental evidence, functional annotation, and predictive value. Feature set FS2300 was included in one of the top-performing validated models, achieving up to a 13% F1-score improvement over baseline. It was the only set that consistently generalized across all tested algorithms (SVM, XGBoost, FNN, Ensemble). Notably, it included a gene encoding for a predicted glutathione-gated potassium efflux system protein (*kefB*, BCM300 RS02305). This efflux system helps maintain intracellular K^+^ homeostasis and belongs to the monovalent cation:proton antiporter (CPA2) family.^22–24^ In *H. pylori*, the expression of *kefB* has been shown to increase, following exposure to metronidazole *in vitro* and to be higher in multidrug-resistant clinical strains compared to susceptible strains.^25^ Based on the importance score, two features contributed highly to the predictive performance of this model: 459267 C (cytosine at position 459267 bp associated with resistance) and RS02305 93V (valine at codon 93 associated with resistance). The 93V mutation is located in a transmembrane domain (TM4) and could theoretically influence the structure of the channel, potentially affecting efflux pump activity. The 459267 C intergenic mutation is located in the initially transcribed region of the 5 ‘UTR of *kefB*, only 10 bp downstream of the transcription start site, and thus could influence its transcription via secondary structures.^26^

Interestingly, three feature sets contained genes related to membrane biogenesis, which may influence metronidazole permeability. In FS4140, *lolF* (BCM300 RS04145) and *secA* (BCM300 RS04140) encode proteins related to lipoprotein release and protein export through the inner membrane, respectively.^27–29^ FS4140 and FS2570 also included outer membrane proteins (OMPs) *hofF* (BCM300 RS04150) and *hofC* (BCM300 RS02570), both members of the Hof family. While their functions remain unclear, *hofF* has been linked to gastric cell adhesion^30^, and *hofC* has been shown to be essential for gastric colonization in a murine model.^31,32^ FS4435 contained *fadL* (BCM300 RS04455), an ortholog of the *E. coli* gene involved in fatty acid transport and membrane integrity.^33^ Altogether, mutations affecting these genes may alter membrane properties and the diffusion of metronidazole.

Additionally, FS7650 included a zinc-metallopeptidase (BCM RS07655), part of the M48 family involved in membrane protein quality control.^34,35^ FS4435 also featured *hup* (BCM300 RS04435), a histone-like DNA-binding protein that, among other predicted functions, protects against oxidative stress–induced DNA damage in *H. pylori*.^36^ A variant 28 bp upstream of *hup* in the 5 UTR may affect its expression and enhance resistance to metronidazole.

## 3 Discussion

Antibiotic resistance is one of the primary reasons responsible for the failure of treatment against *H. pylori*.^1,37^ Since cancer rates subsequent to *H. pylori* infections are increasing on a global scale^4^ improving treatment and predicting treatment failures are key priorities. In many patients, *H. pylori* is resistant to at least one antibiotic included in recommended therapies, with resistance rates exceeding 15% in most regions.^7^ While phenotypic susceptibility testing remains the gold standard in diagnostic laboratories and is currently recommended prior to *H. pylori* eradication by recent clinical guidelines (Maastricht VI/Florence report)^6^, its implementation is limited by the fastidious growth requirements of *H. pylori*. Consequently, sequence-based testing represents a faster and more practical culture-independent alternative for susceptibility-guided therapy in clinical settings. However, established sequence-based assays are typically non-algorithmic and only consider the presence, absence or variation of one or few out of a limited set of sequence markers. Here, we used supervised ML to build and apply predictive models for *H. pylori* antibiotic resistance using a dataset generated from 500 clinical isolates. To our knowledge, this represents the largest dataset for *H. pylori* that combines whole-genome sequences with phenotypic susceptibility testing for clarithromycin, levofloxacin, and metronidazole.

To establish a baseline for evaluating ML performance, we performed (non-algorithmic) predictions with the same genetic variants used as susceptibility markers in established sequence-based assays for *H. pylori*. We could confirm high predictive performance of these genetic markers for clarithromycin and levofloxacin, which is in agreement with previous studies^19,38^ and commercially available assays.^15–18^ In contrast, sequence-based predictions for metronidazole resistance were notably less accurate, with a sensitivity below 50%. These findings are consistent with previous report of poor concordance between sequence-based analysis and phenotypic testing for metronidazole resistance.^14,19,39^ By applying ML, we were able to improve resistance prediction by integrating less-characterized genetic variants in addition to the high-confidence resistance markers used in molecular assays. Unlike the non-algorithmic univariate approach, ML models displayed small to large increases in predicted sensitivity for all tested antibiotics, particularly with the FNN classifier, although overall performance for metronidazole remained suboptimal, with a sensitivity below 70%. Compared to single locus-based assays, the ML models would thus produce fewer false-negatives, indicating that the ML approach was already able to capture additional genotype-to-phenotype relationships related to antibiotic resistance. In a clinical setting, a higher true-positive rate would reduce the likelihood of prescribing ineffective treatments, therefore improving the success of eradication therapy and contributing to more effective antibiotic stewardship. One current limitation of this approach is that high-quality whole-genome sequencing still requires DNA from cultured isolates and remains relatively costly. However, sequencing costs continue to decline, and obtaining *H. pylori* genome sequences directly from biopsy material without prior culture is likely to become feasible and cost-effective in the near future.

To further improve predictive performance for metronidazole resistance from whole genome data, we developed a feature selection pipeline that combined sliding-window evaluation with Shapley-based importance scoring to identify additional candidate genotypes. This approach yielded a group of 32 feature sets that were used for subsequent model training. Based on F1-score improvements, 11 individual sets and 53 synergistic combinations were selected for further validation. Empirical testing of the selected models with an independent cohort of 50 *H. pylori* clinical isolates confirmed the generalizability of a total of 48 models. Considering the high genetic variability of *H. pylori*, a model’s ability to generalize to new data is essential for maintaining consistent performance across isolates. Using only the known genetic markers for resistance to metronidazole, we observed a sharp drop in performance between the training (HPRES-500) and validation (HPRES-50) cohorts. Using similar genetic markers, previous studies based on different cohorts also reported variable results^14,19,39^, further confirming the limited reliability of current genetic markers. Since the cohorts used so far to evaluate metronidazole resistance (including ours) originate from geographically diverse regions of the globe (Europe, USA, Central Africa, Southeast Asia), a potential influence of *H. pylori* phylogeographic diversity on model performance cannot be excluded. While some inter-cohort variation was also observable with ML models, the relative F1-score improvement over baseline was, on average, higher in the training cohort than in the validation cohort. Our results suggest that our whole-genome ML approach provides the greatest benefit in isolates where classical molecular assays struggle the most to predict metronidazole resistance.

The best-performing ML model in our study achieved accuracies of approximately 75% across all tested cohorts, indicating that it could be suitable for metronidazole susceptibility prediction in a clinical setting. Previous studies have also applied ML approaches to predict antibiotic resistance in *H. pylori*. One study^40^ reported close to 100% accuracy for metronidazole resistance prediction based on a single genotype in *addA* (HP1553), a gene encoding a helicase-nuclease involved in DNA recombinational repair.^41^ However, this finding was based on a small cohort of only 49 *H. pylori* isolates from China, with a strong class imbalance (92% of resistant samples), raising concerns about its general applicability in more diverse sample groups. Another recent study^42^ used XGboost and 10 different genetic variants to reach 84% accuracy. The top-ranked features in this model were located in a type II methyltransferase (*mod*, HP0260), a predicted transcriptional regulator (*hypF*, HP0048), and a hypothetical protein (HP0080). Notably, none of these loci overlapped with the genetic variants identified in our study. Although the dataset used in this work was larger (296 isolates), it still showed moderate class imbalance with a 69% resistance rate for metronidazole. Because the training cohort mostly consisted of *H. pylori* isolates from Asia, and model generalization was not empirically tested on an independent validation cohort, a direct comparison with our model is limited.

Functional interpretation of our most predictive features revealed plausible biological mechanisms. In addition to revealing which genetic variants contributed the most to prognostic performance in known genetic markers such as *rdxA* and *frxA*, several novel candidates were identified. In particular, the glutathione-gated potassium efflux system protein *kefB* (HP0471) was part of generalizable ML models and has previously been shown to be upregulated in metronidazole-resistant strains.^25^ In *H. pylori*, potassium homeostasis has been demonstrated to be essential for colonization in a murine model^43^, and the transcription of *kefB* was shown to be regulated by the stringent response regulator SpoT.^44^ The exact mechanism connecting KefB to metronidazole resistance has not been described. It does not belong to efflux pump superfamilies which are known to be associated with antimicrobial resistance, such as ABC, MATE, or RND.^45^ The increased expression of *kefB*, correlating with metronidazole resistance, was previously associated with a higher efflux activity and biofilm formation ^25^. These traits may be directly responsible for resistance by increasing drug efflux and decreasing drug uptake.^46^ The *kefB* overexpression was also shown to be co-dependent with the expression of two other efflux pumps, which could subsequently increase drug efflux even more.^25^ Furthermore, a K+ efflux can modify membrane potential and could decrease diffusion of metronidazole through the membrane. Because potassium homeostasis is important for numerous cellular processes^47,48^, KefB might also influence other mechanisms involved in metronidazole resistance indirectly, such as prodrug activation or DNA damage repair. Furthermore, we propose several synergistic models combining genetic variants from genes associated with distinct functional categories (such as efflux, transport, and DNA damage mitigation) further supporting the hypothesis that resistance to metronidazole can be a complex and multifactorial trait.

In conclusion, this study demonstrates that ML approaches can improve antibiotic resistance prediction in *H. pylori* by overcoming the limitations of traditional nucleotide sequence-based diagnostic assays, particularly for antibacterials which can have complex and polygenic resistance mechanisms. The integration of whole-genome data with ML methods not only improved predictive performances but also highlighted novel genetic determinants of resistance to metronidazole that can be functionally characterized in future experimental studies. Further validation of the ML models trained in this study in additional cohorts and integration into comprehensive diagnostic platforms can, as enhancements of phenotypic testing, help instruct susceptibility-guided therapies for more effective treatment of *H. pylori* infection.

## 4 Material and Methods

### 4.1 Strain Collection

A total of 533 strains of *H. pylori* were included in the collection (Table S1), 483 as the discovery cohort and 50 as an independent test cohort. Strains were routinely cultured on blood agar plates (blood agar base II; Oxoid, Germany) containing 10% horse blood, 10 mg/l vancomycin, 3.2 mg/l polymyxin B, 4 mg/l amphotericin B and 5 mg/l trimethoprim. Incubation was performed at 37°C under microaerobic conditions using Anaerocult C gas generating bags (Merck, Germany) in air-tight jars (Oxoid, Germany).

### 4.2 Antibiotic susceptibility testing

Antibiotic susceptibility was determined for clarithromycin, levofloxacin, and metronidazole using E-test strips (bioMérieux, France). EUCAST clinical breakpoint values were used to interpret E-test results (Version 8.1, 2018. http://www.eucast.org). For clarithromycin, a Minimum Inhibitory Concentration (MIC) below or equal to 0.25 mg/L was considered susceptible, and above 0.25 mg/L was considered resistant. For levofloxacin, an MIC below or equal to 1 mg/L was considered susceptible, and above 1 mg/L was considered resistant. For metronidazole, an MIC below or equal to 8 mg/L was considered susceptible, and above 8 mg/L was considered resistant.

### 4.3 Genome sequencing

Genomic DNA was extracted using genomic-tip 20/G anion-exchange columns according to the manufacturer’s protocol (Qiagen, Germany). All samples were prepared using the Nextera XT library preparation kit and sequenced with >30-fold coverage on an Illumina MiSeq instrument with 600-cycle v3 reagent cartridges. Library quality control was performed using a TapeStation 4200 system with the High Sensitivity D5000 kit (Agilent Technologies, USA) and a Qubit fluorometer with the High Sensitivity dsDNA kit (Invitrogen, USA).

### 4.4 Feature extraction and encoding

Sequencing reads were aligned to the *H. pylori* reference genome BCM300 (RefSeq accession NZ LT837687.1) using BWA version 0.7.17 with the BWA-MEM algorithm.^49^ Variant calling and effect prediction were performed using VarScan 2.4.2^50^, snpEff 4.3^51^, and Geneious Prime 2022.1.1 (https://www.geneious.com). Variants were classified into three distinct feature categories:

- (A) Intergenic and non-coding variants, such as rRNAs;
- (B) Non-synonymous variants resulting in amino-acid changes;
- (C) Insertions or deletions causing frameshifts, or premature stop codons leading to truncated proteins.

For downstream ML, variants were encoded as binary features. Category A features represented the presence of a specific nucleotide at a given position (e.g., 1458 A for adenine at position 1458). Category B features indicated the presence of a specific amino acid at a codon within a gene (e.g., RS03700 87 N for asparagine at codon 87 of gene RS03700). Category C features encoded gene status as either intact or truncated. Each feature can take a value of 1 or 0, indicating the presence or absence of a nucleotide or amino-acid for categories A and B or the truncated or complete form of a gene for category C.

### 4.5 Feature partitioning for predictive power evaluation

A sliding-window approach was used to reduce feature dimensionality and computational burden by grouping features according to genomic location. In this step, the genome is partitioned into non-overlapping windows based on the base-pair (bp) locations as illustrated in Fig. 3A. This approach reduces the computational load by learning with a more manageable number of features within each window instead of dealing with hundreds of thousands of features simultaneously. Here, we chose a window size of 2000 base-pair (bps), because among all windows with this length, the median number of features is around 600, which is not too overwhelming for the ML models, but still covers a diverse range of features. Different window sizes can be experimented with to optimize the balance between local detail and computational efficiency. This results in around 830 non-overlapping windows, each encapsulating a subset of mutation features with all categories of features, i.e., CatA, CatB, and CatC.

**Fig. 3.**
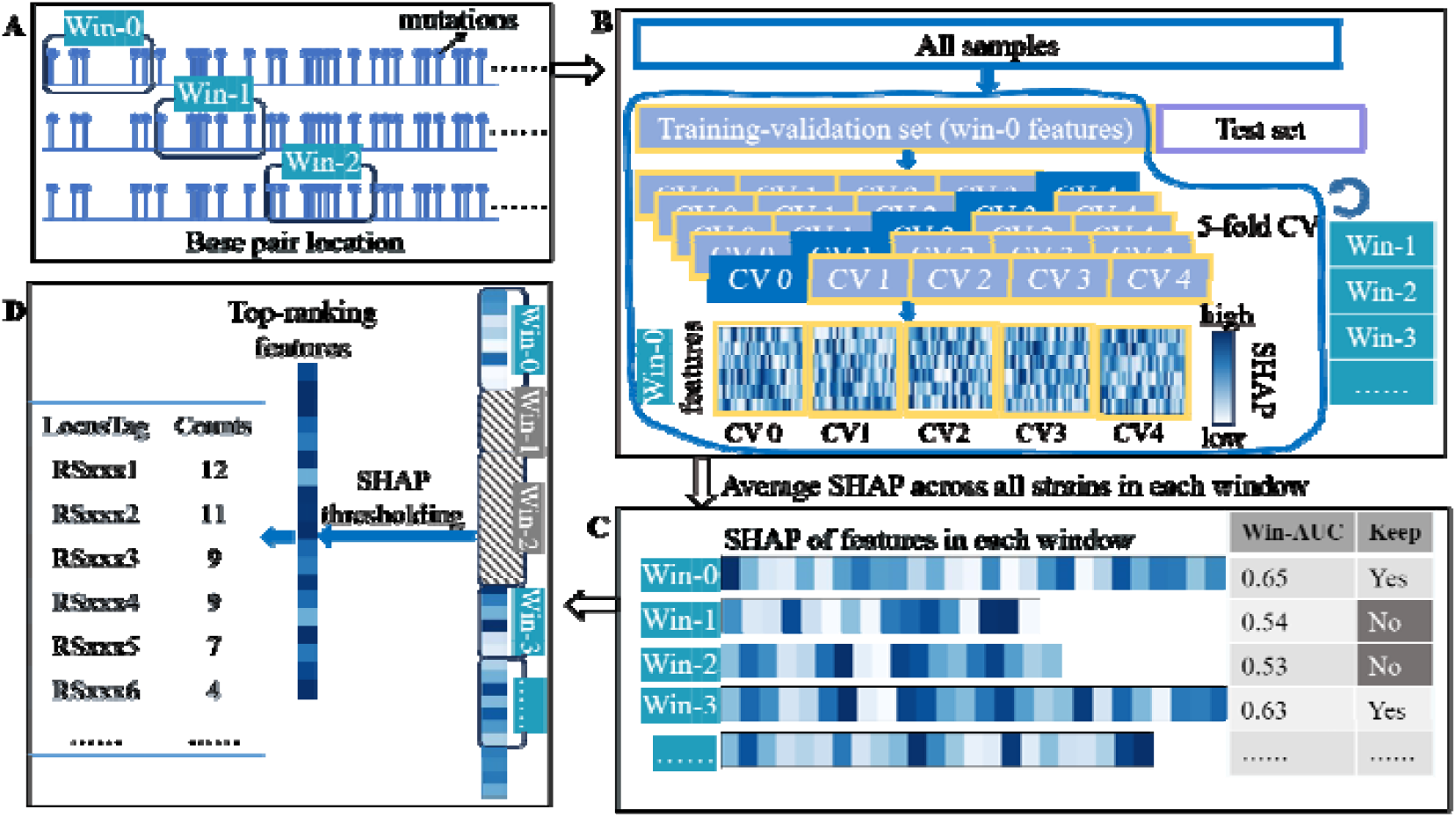
Schematic of sliding window feature selection. **A:** Sliding window feature partitioning, with each window having 2000 base pairs. Note that the number of features might be different in each window. **B:** After the (80:20) train-test split of all the samples in the HPRES-500 dataset, feature selection is done only with the **training** set. Feature SHAP scores are computed in each fold for each window. Notably, the number of features in each window is not necessarily the same. **C:** ROC-AUC is computed for each window, as well as the averaged SHAP scores for all features in each window. Those windows with an AUC (“Win-AUC”) above a threshold, here set to 0.6, are kept for subsequent analysis. **D:** The averaged SHAP scores of the remaining windows are concatenated and go through thresholding, and the occurrence count of LocusTags that passed the threshold is calculated. CV: cross-validation. AUC: area under the Receiver Operating Characteristics curve.

**Fig. 4.**
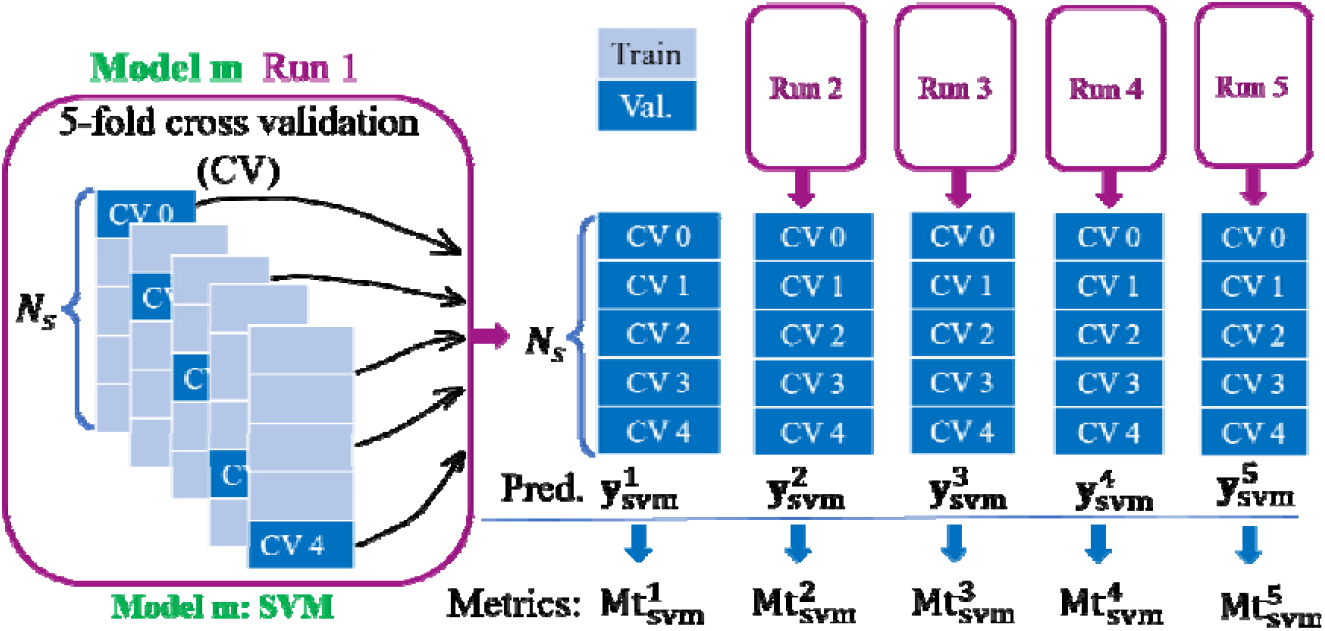
Schematic of multiple runs of five-fold cross-validation for model *m*. In a 5-fold cross-validation, the dataset is divided into five subsets. Each subset is used once as the test set, while the remaining subsets are used for training. After each run, the prediction for the whole dataset is collected, denoted as 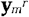 and model *m* is set to SVM for illustration. The total number of samples are denoted as *N*_*s*_. This collected prediction is subsequently used to compute the performance metrics 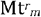. Furthermore, multiple runs ensure the robustness and reliability of the model performance metrics.

### 4.6 Region-specific feature evaluation using ML

To ensure that there was no data leakage in feature selection, we first split our entire HpRES-500 dataset into a training set and a withheld test set with a ratio of 80:20. Then the feature selection was exclusively conducted on the training set, illustrated in Fig 3B. Within each window, we trained an XGBoost model to assess the predictive capabilities and feature importance within that specific genomic region with multiple independent runs of five-fold cross-validation. This localized analysis enabled us to identify potential sequence-based markers and gain insights into the significance of multiple mutations across different regions of the genome.

The averaged ROC-AUC was computed across all five folds for each window, and those with an average ROC-AUC over 0.55 were included in our subsequent SHAP score thresholding analysis. This led to the inclusion of more than 36,000 features for metronidazole from the top-performing windows. These features then formed the feature pool for the subsequent further selection.

### 4.7 Shapley analysis for feature selection

Shapley values were used to quantify the contribution of individual features within each genomic window selected during model training. Values were computed per sample and per feature using the SHAP framework^21^, which estimates the average marginal contribution of a feature across all possible combinations with other features, accounting for feature interactions and dependencies. SHAP unifies several interpretability methods, including Shapley sampling^52^, Shapley regression, LIME^53^, DeepLIFT^54^, layer-wise relevance propagation^55^, and QII.^56^

An average SHAP value (noted as the ‘feature importance score’) was calculated across all cross-validation folds obtained during training for every feature within the top-performing genomic windows (avg. ROC-AUC > 0.55). For example, when a feature is ON (feature value = 1, indicating the presence of a given nucleotide for category A, an amino-acid for category B, or an inactive protein for category C), an average SHAP score of 0.1 for antibiotic susceptibility (class 1) and −0.1 for antibiotic resistance (class 0) indicates that this feature contributes 0.1 toward the prediction of antibiotic susceptibility. Conversely, when this same feature is OFF (feature value = 0, indicating the absence of the nucleotide or amino-acid, or an active protein), it would contribute to the prediction of antibiotic resistance. A threshold of 0.01 for the feature importance score was chosen to balance feature selection and model performance.

### 4.8 Frequency of important features within genes

Features that passed the filtering thresholds from (i) the sliding-window ROC-AUC analysis and (ii) feature importance score analysis were aggregated at the gene level on their genomic coordinates. Features mapping to the same locus or in close proximity (within 10,000 bp) were grouped as one feature set, resulting in a total of 32 feature sets being identified for the metronidazole labelled dataset.

### 4.9 Model training and validation

Three ML models were evaluated for binary classification of antibiotic resistance (class 0: resistant, class 1: susceptible): XGBoost, Support Vector Machine (SVM), and a Fully Connected Neural Network (FNN). XGBoost, an ensemble gradient boosting algorithm, was optimized using Bayesian search via the Optuna framework ^57^ to tune hyperparameters such as learning rate, tree depth, and number of estimators. SVMs employed non-linear kernels, with regularization parameters and kernel coefficients also tuned using Bayesian optimization with Optuna.^57^

The FNN consisted of three dense layers with 128, 32, and 2 units, using ReLU activation functions, batch normalization, and a dropout rate of 0.58 to improve generalization. The final layer used a Softmax activation function for classification. The network was implemented in TensorFlow^58^ and trained using the Adam optimizer with binary cross-entropy loss and a learning rate of 0.001. An EarlyStopping callback was used to reduce overfitting. All models were trained with each feature sets identified by the selection pipeline.

Robust performance estimates were obtained by conducting NR independent runs of 5-fold cross-validation for each model. Predictions from all folds and runs were aggregated to calculate classification metrics, including accuracy, precision, recall, F1 score, and area under the receiver operating characteristic curve (AUC). Models were evaluated both independently and as part of an ensemble combining predictions from all models.

#### 4.9.1 Single Model Evaluation

In this evaluation scheme, multiple runs of five-fold cross-validation were performed and the performance metrics were computed for each model in each run separately. In each cross-validation fold, test fold predictions are collected, and concatenating these across all five folds in one run yields the predictions for all samples for that run, denoted as 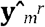, where *m* is the model name and *r* is the run index of the run. In this evaluation scheme, the performance metrics are computed with the collected prediction of all samples, 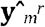, and their ground truth **y**^*r*^ for each model in each run separately, denoted as 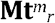.

The average performance for model *m*, then is given by

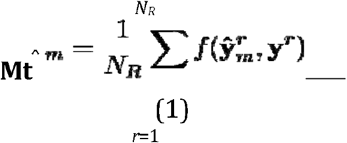

where 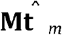 is the performance averaged over all runs for model *m*. *N*_*R*_ is the total number of runs, and the function *f*(·) takes the prediction 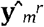 and the ground truth y as input to compute all averaged performance metrics, denoted “Avr.” in the text.

#### 4.9.2 Aggregation Across Multiple Runs Within Each Model

This method averages predictions across multiple runs for each model and computes performance metrics once. For a given model, the concatenated predictions from five validation folds of all runs for this model are 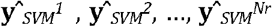, where N is the total umber of runs. These predictions are firstly averaged to get an overall prediction from this model, i.e., 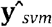, given by

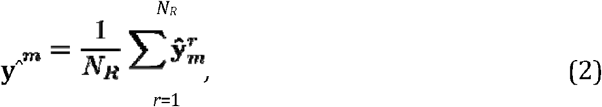

Subsequently, the averaged prediction 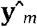 and the ground truth y are used as input to *f*(·) to compute aggregated performance metrics, denoted “Agg.” in the text.

#### 4.9.3 Ensemble Across Multiple Models

Ensemble models combine predictions from multiple ML algorithms to improve accuracy, robustness, and generalization.^59^ This aims to aggregate predictions from multiple models across multiple runs. Compared to the aggregation method applied within each model, ensemble models further integrate predictions from other models for the same sample.

To this end, we implemented a stacking ensemble, where the concatenated predictions from all validation folds, specifically, 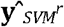 from the 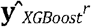 from XGBoost, and 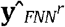 from the FNN at run *r*, were firstly aggregated using a linear averaging operation, denoted “Ensemble” in the text. Subsequently, model performances were evaluated using two schemes: (1) aggregation across multiple runs of a single model (“Agg.”), and (2) aggregation across multiple runs and models (“Ensemble”). The prediction calculation of this ensemble models is given by

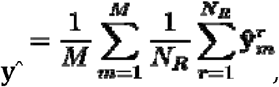

where y^ is the prediction of the ensemble model averaged over *N*_R_ runs, over *M* models, and 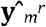 is the prediction from the *r*-th run for model *m*. Subsequently, y^ is used to compute the performance metrics such as AUC, accuracy, sensitivity, specificity, etc.

## 5. Supplementary material

- Supplementary Table S1. Summary of *H. pylori* strains included in the training and validation datasets, and corresponding antibiotic resistance profiles.
- Supplementary Table S2. Known biomarkers associated with antibiotic resistance.
- Supplementary Table S3. Genomic clusters identified after feature selection for metronidazole resistance prediction, including matching functional categories.
- Supplementary Table S4. F1-score performance of individual and grouped feature sets in training and validation datasets for metronidazole resistance prediction.
- Supplementary Table S5. Functional assignment of feature sets with statistically significant improvement over baseline performance in metronidazole resistance prediction.
- Supplementary Table S6. Generalizable feature sets achieving an F1-score ≥ 0.65 in both training and validation cohorts for metronidazole resistance prediction.
- Supplementary Table S7. Complete performance reports for generalizable feature sets used in metronidazole resistance prediction.

## 6 Data availability

Raw sequencing data generated in this study were deposited in the National Center for Biotechnology Information Sequence Read Archive database under BioProject PRJNA1295820.

## 7 Code Accessibility

Source code can be access at https://github.com/DiyuanLu/HelicoPredict-new#

8 Acknowledgments

The authors would like to thank Gudrun Pfaffinger, Evelyn Weiss, Christopher Beck, Cornelia Fük, Martina Merker, Elke Stummeyer, Teresa Burrell and Carolina Domingos for technical assistance with strain culture, antibiotic susceptibility testing, and genome sequencing.

## 10 Ethics statement

The samples included in this project were provided by multiple studies, individually approved by local ethics committee and government authorities and was conducted in accordance with current Good Clinical Practice guidelines and the Declaration of Helsinki (https://doi.org/10.1001/jama.2013.281053).

## 12 Responsible Use of Artificial Intelligence

Artificial intelligence (AI) tools including ChatGPT and Grammarly were employed in this manuscript to refine grammar, spelling, and readability. These tools were used exclusively for linguistic enhancement, ensuring clarity and coherence without altering the scientific content, interpretation, or conclusions. All intellectual contributions, data analyses, and critical discussions remain the work of the authors. The final manuscript was thoroughly reviewed and approved by all co-authors to ensure accuracy and integrity.

## References

1. Malfertheiner, P., Camargo, M.C., El-Omar, E., Liou, J.M., Peek, R., Schulz, C., Smith, S.I., and Suerbaum, S. (2023). Helicobacter pylori infection. Nat Rev Dis Primers 9, 19.

2. Liou, J.-M., Malfertheiner, P., Lee, Y.-C., Sheu, B.-S., Sugano, K., Cheng, H.-C., Yeoh, K.-G., Hsu, P.-I., Goh, K.-L., and Mahachai, V. (2020). Screening and eradication of Helicobacter pylori for gastric cancer prevention: the Taipei global consensus. Gut 69, 2093–2112.

3. Chiang, T.-H., Chang, W.-J., Chen, S.L.-S., Yen, A.M.-F., Fann, J.C.-Y., Chiu, S.Y.-H., Chen, Y.-R., Chuang, S.-L., Shieh, C.-F., and Liu, C.-Y. (2021). Mass eradication of Helicobacter pylori to reduce gastric cancer incidence and mortality: a long-term cohort study on Matsu Islands. Gut 70, 243–250.

4. Park, J.Y., Georges, D., Alberts, C.J., Bray, F., Clifford, G., and Baussano, I. (2025). Global lifetime estimates of expected and preventable gastric cancers across 185 countries. Nature Medicine, 1–8.

5. Lee, Y.-C., Chiang, T.-H., Chou, C.-K., Tu, Y.-K., Liao, W.-C., Wu, M.-S., and Graham, D.Y. (2016). Association between Helicobacter pylori eradication and gastric cancer incidence: A systematic review and meta-analysis. Gastroenterology 150, 1113–1124.

6. Malfertheiner, P., Megraud, F., Rokkas, T., Gisbert, J.P., Liou, J.-M., Schulz, C., Gasbarrini, A., Hunt, R.H., Leja, M., and O’Morain, C. (2022). Management of Helicobacter pylori infection: the Maastricht VI/Florence consensus report. Gut 71, 1724–1762.

7. Savoldi, A., Carrara, E., Graham, D.Y., Conti, M., and Tacconelli, E. (2018). Prevalence of antibiotic resistance in Helicobacter pylori: a systematic review and meta-analysis in World Health Organization regions. Gastroenterology 155, 1372-1382. e1317.

8. Schulz, C., Liou, J.-M., Alboraie, M., Bornschein, J., Nunez, C.C., Coelho, L.G., Quach, D.T., Fallone, C.A., Chen, Y.-C., and Gerhard, M. (2025). Helicobacter pylori antibiotic resistance: a global challenge in search of solutions. Gut.

9. Lin, C.-H., Hsu, P.-I., Tseng, C.-D., Chao, P.-J., Wu, I.T., Ghose, S., Shih, C.-A., Lee, S.-H., Ren, J.-H., Shie, C.-B., and et al. (2023). Application of artificial intelligence in endoscopic image analysis for the diagnosis of a gastric cancer pathogen Helicobacter pylori infection. Scientific Reports 13, 13380.

10. Itoh, T., Kawahira, H., Nakashima, H., and Yata, N. (2018). Deep learning analyzes Helicobacter pylori infection by upper gastrointestinal endoscopy images. Endoscopy international open 6, E139–E144.

11. Zhou, S., Marklund, H., Blaha, O., Desai, M., Martin, B., Bingham, D., Berry, G.J., Gomulia, E., Ng, A.Y., and Shen, J. (2020). Deep learning assistance for the histopathologic diagnosis of Helicobacter pylori. Intelligence-Based Medicine 1, 100004.

12. Klein, S., Gildenblat, J., Ihle, M.A., Merkelbach-Bruse, S., Noh, K.-W., Peifer, M., Quaas, A., and Bttner, R. (2020). Deep learning for sensitive detection of Helicobacter pylori in gastric biopsies. BMC gastroenterology 20, 1–11.

13. Ailloud, F., Estibariz, I., and Suerbaum, S. (2021). Evolved to vary: genome and epigenome variation in the human pathogen Helicobacter pylori. FEMS Microbiology Reviews 45, fuaa042.

14. Tshibangu-Kabamba, E., and Yamaoka, Y. (2021). Helicobacter pylori infection and antibiotic resistance—from biology to clinical implications. Nature Reviews Gastroenterology & Hepatology 18, 613–629.

15. Cambau, E., Allerheiligen, V., Coulon, C., Corbel, C., Lascols, C., Deforges, L., Soussy, C.J., Delchier, J.C., and Megraud, F. (2009). Evaluation of a new test, genotype HelicoDR, for molecular detection of antibiotic resistance in Helicobacter pylori. J Clin Microbiol 47, 3600–3607.

16. Jehanne, Q., Benejat, L., Megraud, F., Bessede, E., and Lehours, P. (2020). Evaluation of the Allplex H. pylori and ClariR PCR assay for Helicobacter pylori detection on gastric biopsies. Helicobacter 25, e12702.

17. Lottspeich, C., Schwarzer, A., Panthel, K., Koletzko, S., and Russmann, H. (2007). Evaluation of the novel Helicobacter pylori ClariRes real-time PCR assay for detection and clarithromycin susceptibility testing of H. pylori in stool specimens from symptomatic children. J Clin Microbiol 45, 1718–1722.

18. Schuetz, A.N., Theel, E.S., Cole, N.C., Rothstein, T.E., Gordy, G.G., and Patel, R. (2024). Testing for Helicobacter pylori in an era of antimicrobial resistance. J Clin Microbiol 62, e0073223.

19. Hulten, K.G., Genta, R.M., Kalfus, I.N., Zhou, Y., Zhang, H., and Graham, D.Y. (2021). Comparison of culture with antibiogram to Next-Generation Sequencing using bacterial isolates and formalin-fixed, paraffin-embedded gastric biopsies. Gastroenterology 161, 1433–1442 e1432.

20. Goodwin, A., Kersulyte, D., Sisson, G., Veldhuyzen van Zanten, S.J., Berg, D.E., and Hoffman, P.S. (1998). Metronidazole resistance in Helicobacter pylori is due to null mutations in a gene (rdxA) that encodes an oxygen-insensitive NADPH nitroreductase. Molecular microbiology 28, 383–393.

21. Lundberg, S.M., and Lee, S.-I. (2017). A unified approach to interpreting model predictions. Advances in neural information processing systems 30.

22. Ferguson, G.P., Munro, A.W., Douglas, R.M., McLaggan, D., and Booth, I.R. (1993). Activation of potassium channels during metabolite detoxification in Escherichia coli. Molecular microbiology 9, 1297–1303.

23. Rasmussen, T. (2023). The potassium efflux system Kef: bacterial protection against toxic electrophilic compounds. Membranes 13, 465.

24. Saier, M.H., Jr., Eng, B.H., Fard, S., Garg, J., Haggerty, D.A., Hutchinson, W.J., Jack, D.L., Lai, E.C., Liu, H.J., Nusinew, D.P., et al. (1999). Phylogenetic characterization of novel transport protein families revealed by genome analyses. Biochim Biophys Acta 1422, 1–56.

25. Cai, Y., Wang, C., Chen, Z., Xu, Z., Li, H., Li, W., and Sun, Y. (2020). Transporters HP0939, HP0497, and HP0471 participate in intrinsic multidrug resistance and biofilm formation in Helicobacter pylori by enhancing drug efflux. Helicobacter 25, e12715.

26. Laalami, S., Zig, L., and Putzer, H. (2014). Initiation of mRNA decay in bacteria. Cell Mol Life Sci 71, 1799–1828.

27. McClain, M.S., Bryant, K.N., McDonald, W.H., Algood, H.M.S., and Cover, T.L. (2023). Identification of an essential LolD-like protein in Helicobacter pylori. Journal of Bacteriology 205, e00052.-00023.

28. Liechti, G., and Goldberg, J.B. (2012). Outer membrane biogenesis in Escherichia coli, Neisseria meningitidis, and Helicobacter pylori: Paradigm deviations in H. pylori. Front Cell Infect Microbiol 2, 29.

29. Tsirigotaki, A., De Geyter, J., Šoštaric’, N., Economou, A., and Karamanou, S. (2017). Protein export through the bacterial Sec pathway. Nature Reviews Microbiology 15, 21–36.

30. Ji, X., Wang, Y., Li, J., Rong, Q., Chen, X., Zhang, Y., Liu, X., Li, B., and Zhao, H. (2017). Application of FLP-FRT system to construct unmarked deletion in Helicobacter pylori and functional study of gene hp0788 in pathogenesis. Front Microbiol 8, 2357.

31. Kavermann, H., Burns, B.P., Angermuller, K., Odenbreit, S., Fischer, W., Melchers, K., and Haas, R. (2003). Identification and characterization of Helicobacter pylori genes essential for gastric colonization. The Journal of experimental medicine 197, 813–822.

32. Baldwin, D.N., Shepherd, B., Kraemer, P., Hall, M.K., Sycuro, L.K., Pinto-Santini, D.M., and Salama, N.R. (2007). Identification of Helicobacter pylori genes that contribute to stomach colonization. Infection and immunity 75, 1005–1016.

33. Tan, Z., Black, W., Yoon, J.M., Shanks, J.V., and Jarboe, L.R. (2017). Improving Escherichia coli membrane integrity and fatty acid production by expression tuning of FadL and OmpF. Microb Cell Fact 16, 38.

34. Akiyama, Y. (2009). Quality control of cytoplasmic membrane proteins in Escherichia coli. J Biochem 146, 449–454.

35. Sakoh, M., Ito, K., and Akiyama, Y. (2005). Proteolytic activity of HtpX, a membrane-bound and stress-controlled protease from Escherichia coli. J Biol Chem 280, 33305–33310.

36. Wang, G., Lo, L.F., and Maier, R.J. (2012). A histone-like protein of Helicobacter pylori protects DNA from stress damage and aids host colonization. DNA Repair (Amst) 11, 733–740.

37. Nezami, B.G., Jani, M., Alouani, D., Rhoads, D.D., and Sadri, N. (2019). Helicobacter pylori mutations detected by Next-Generation Sequencing in formalin-fixed, paraffin-embedded gastric biopsy specimens are associated with treatment failure. J Clin Microbiol 57.

38. Chen, M.J., Chen, P.Y., Fang, Y.J., Bair, M.J., Chen, C.C., Chen, C.C., Yang, T.H., Lee, J.Y., Yu, C.C., Kuo, C.C., et al. (2023). Molecular testing-guided therapy versus susceptibility testing-guided therapy in first-line and third-line Helicobacter pylori eradication: Two multicentre, open-label, randomised controlled, non-inferiority trials. Lancet Gastroenterol Hepatol 8, 623–634.

39. Tuan, V.P., Narith, D., Tshibangu-Kabamba, E., Dung, H.D.Q., Viet, P.T., Sokomoth, S., Binh, T.T., Sokhem, S., Tri, T.D., Ngov, S., et al. (2019). A Next-Generation Sequencing-based approach to identify genetic determinants of antibiotic resistance in Cambodian Helicobacter pylori clinical isolates. J Clin Med 8.

40. Yu, J., Jia, Y., Yu, Q., Lin, L., Li, C., Chen, B., Zhong, P., Lin, X., Li, H., Sun, Y., et al. (2023). Deciphering complex antibiotic resistance patterns in Helicobacter pylori through whole genome sequencing and machine learning. Front Cell Infect Microbiol 13, 1306368.

41. Amundsen, S.K., Fero, J., Hansen, L.M., Cromie, G.A., Solnick, J.V., Smith, G.R., and Salama, N.R. (2008). Helicobacter pylori AddAB helicase-nuclease and RecA promote recombination-related DNA repair and survival during stomach colonization. Mol Microbiol 69, 994–1007.

42. Wang, Y., Shuwen, Z., Rui, G., Yanke, L., Honghao, Y., Xunan, Q., Jijun, C., Chuxuan, N., Yuan, Y., and and Gong, Y. (2025). Assessment for antibiotic resistance in Helicobacter pylori: A practical and interpretable machine learning model based on genome-wide genetic variation. Virulence 16, 2481503.

43. Stingl, K., Brandt, S., Uhlemann, E.M., Schmid, R., Altendorf, K., Zeilinger, C., Ecobichon, C., Labigne, A., Bakker, E.P., and de Reuse, H. (2007). Channel-mediated potassium uptake in Helicobacter pylori is essential for gastric colonization. Embo J 26, 232–241.

44. Geng, X., Li, W., Chen, Z., Gao, S., Hong, W., Ge, X., Hou, G., Hu, Z., Zhou, Y., Zeng, B., et al. (2017). The bifunctional enzyme SpoT is involved in the clarithromycin tolerance of Helicobacter pylori by upregulating the transporters HP0939, HP1017, HP0497, and HP0471. Antimicrob Agents Chemother 61.

45. Krzyzek, P. (2024). Helicobacter pylori efflux pumps: A double-edged sword in antibiotic resistance and biofilm formation. Int J Mol Sci 25.

46. Tshibangu-Kabamba, E., Ngoma-Kisoko, P.J., Tuan, V.P., Matsumoto, T., Akada, J., Kido, Y., Tshimpi-Wola, A., Tshiamala-Kashala, P., Ahuka-Mundeke, S., Ngoy, D.M., et al. (2020). Next-Generation Sequencing of the whole bacterial genome for tracking molecular insight into the broad-spectrum antimicrobial resistance of Helicobacter pylori clinical isolates from the Democratic Republic of Congo. Microorganisms 8.

47. Do, E.A., and Gries, C.M. (2021). Beyond homeostasis: Potassium and pathogenesis during bacterial infections. Infect Immun 89, e0076620.

48. Stautz, J., Hellmich, Y., Fuss, M.F., Silberberg, J.M., Devlin, J.R., Stockbridge, R.B., and Hanelt, I. (2021). Molecular mechanisms for bacterial potassium homeostasis. J Mol Biol 433, 166968.

49. Li, H., and Durbin, R. (2010). Fast and accurate long-read alignment with Burrows-Wheeler transform. Bioinformatics 26, 589–595.

50. Koboldt, D.C., Zhang, Q., Larson, D.E., Shen, D., McLellan, M.D., Lin, L., Miller, C.A., Mardis, E.R., Ding, L., and Wilson, R.K. (2012). VarScan 2: somatic mutation and copy number alteration discovery in cancer by exome sequencing. Genome Res 22, 568–576.

51. Cingolani, P., Platts, A., Wang le, L., Coon, M., Nguyen, T., Wang, L., Land, S.J., Lu, X., and Ruden, D.M. (2012). A program for annotating and predicting the effects of single nucleotide polymorphisms, SnpEff : SNPs in the genome of Drosophila melanogaster strain w1118; iso-2; iso-3. Fly (Austin) 6, 80–92.

52. Castro, J., Gómez, D., and Tejada, J. (2009). Polynomial calculation of the Shapley value based on sampling. Computers & operations research 36, 1726–1730.

53. Ribeiro, M.T., Singh, S., and Guestrin, C. (2016). “Why should I trust you?” Explaining the predictions of any classifier. pp. 1135–1144.

54. Shrikumar, A., Greenside, P., and Kundaje, A. (2017). Learning important features through propagating activation differences. (PMlR), pp. 3145–3153.

55. Binder, A., Montavon, G., Lapuschkin, S., Müller, K.-R., and Samek, W. (2016). Layer-wise relevance propagation for neural networks with local renormalization layers. (Springer), pp. 63–71.

56. Datta, A., Sen, S., and Zick, Y. (2016). Algorithmic transparency via quantitative input influence: Theory and experiments with learning systems. (IEEE), pp. 598–617.

57. Akiba, T., Sano, S., Yanase, T., Ohta, T., and Koyama, M. (2019). Optuna: A next-generation hyperparameter optimization framework. pp. 2623–2631.

58. Abadi, M., Barham, P., Chen, J., Chen, Z., Davis, A., Dean, J., Devin, M., Ghemawat, S., Irving, G., and Isard, M. (2016). TensorFlow: a system for Large-Scale machine learning. pp. 265–283.

59. Dietterich, T.G. (2000). Ensemble methods in machine learning. (Springer), pp. 1–15.

